# FLASH Radiotherapy Mitigates Radiation-Induced Lymphopenia and Prevents Immunosuppression via Chk1-STAT3 Axis Modulation in a Preclinical Thoracic Irradiation Model

**DOI:** 10.1101/2025.10.21.683770

**Authors:** Rong-Hua Tao, Kevin Liu, Edgardo Aguilar, Min Wang, Sadhna Aggarwal, Denae Neill, Brett Velasquez, Sam Beddar, Albert C. Koong, Radhe Mohan, Emil Schüler, Steven H. Lin

**Author notes:** Rong-Hua Tao and Kevin Liu both contributed equally to this work. Author responsible for statistical analysis (Rong-Hua Tao). Senior Author. **Funding:** Research reported in this publication was supported in part by the P01xxxx, project 2 on lymphopenia, and by UTHealth Innovation for Cancer Prevention Research Training Program Postdoctoral Fellowship (Cancer Prevention and Research Institute of Texas grant RP210042). The content is solely the responsibility of the authors and does not necessarily represent the official views of the National Institutes of Health nor of the Cancer Prevention and Research Institute of Texas. **Conflict of Interest:** Steven H. Lin (SHL) has the following disclosures: Grants: STCube pharmaceuticals, Beyond Spring Pharmaceuticals, Nektar Therapeutics; Advisory Board: AstraZeneca, Creatv Microtech; Consultant: XRAD Therapeutics; Cofounder and Stock options: Seek Diagnostics. **Data Availability:** Research data are stored in an institutional repository and will be shared upon request to the corresponding author.

## Abstract

**Background and Aims:** Radiation-induced lymphopenia (RIL) is a frequent side effect of conventional radiation therapy (CONV RT), due to the high radiosensitivity of circulating lymphocytes. Ultra-high dose rate “FLASH” RT may preferentially spare normal tissue while maintaining tumor control. This study evaluates the impact of single-fraction and multi-fraction thoracic FLASH RT on lymphocyte preservation, apoptosis, and immunosuppressive signaling in mice.

**Methods:** We compared the immunological impact of thoracic FLASH RT and CONV RT in C57BL/6 mice using single-fraction (17 Gy) and multi-fraction (2 Gy × 5) regimens using the Mobetron (IntraOp). Longitudinal blood sampling was performed at multiple time-points post-irradiation through facial vein bleed with flow cytometry analysis for CD4+, CD8+, CD19+, and NK cells to assess lymphocyte counts, apoptotic lymphocytes through Annexin V staining, and immune suppression by examining regulatory T cells (Tregs) and PD-1/PD-L1 expression. Mechanistic studies included immunofluorescence and Western blot analyses of splenic tissues to evaluate Chk1 and STAT3 signaling pathways.

**Results:** In single-fraction RT, FLASH significantly reduced lung and heart fibrosis (p < 0.0001) at 28 weeks post-RT. The FLASH effect was also seen acutely on circulating immune cells, with significantly reduced lymphocyte apoptosis and accelerated recovery of CD4^⁺^, CD8^⁺^, CD3^⁺^, NK, and B cell populations compared to CONV RT in both single-fraction and multi-fraction regimens. Conversely, CONV RT induced long-lasting increases in Tregs and sustained PD-1 and PD-L1 expression on T- and B-cells at 2- and 5-months post-irradiation in both fractionation regimens. Within the spleen, we also found CONV RT induced sustained activation of the Chk1–STAT3 pathway in CD45+ immune cells, which correlates with increased PD-1/PD-L1 expression.

**Conclusion:** FLASH RT mitigates RIL, reduces lymphocyte apoptosis, and prevents long-term immunosuppression by reduced activation of the Chk1–STAT3 pathway. These findings suggest FLASH RT may confer immunological advantages over CONV RT to enhance therapeutic efficacy.

## Introduction

Radiation therapy (RT) is utilized in more than half of patients with cancer with curative and palliative intent(1,2). With increasing technological advances, RT is now increasingly employed as a curative alternative to surgery for early-stage cancers and even for select cases of limited metastatic disease(3). However, the primary factor limiting its curative potential is toxicity to the surrounding healthy tissue, which can restrict the ability to deliver the most effective therapeutic doses(4). Lymphocytes, in particular, are among the most radiosensitive cells in the body and are routinely exposed to unintended radiation while circulating through the tumor vasculature within the radiation field and the surrounding tissue outside of the radiation field, and is thereby one of the key drivers of inflammatory responses to radiation(5).

Conventional dose rate radiation therapy utilized in the clinical setting (CONV RT) often causes severe lymphopenia due to the high radiosensitivity of circulating lymphocytes exposed to even low doses of radiation (<1 Gy) near the tumor site. This radiation-induced lymphopenia (RIL) can be long-lasting, with some patients never regaining normal lymphocyte levels, even years after treatment. Importantly, severe lymphopenia is consistently associated with worse outcomes across nearly all cancer types treated with RT, likely due to impaired immune function(6–10). This immunosuppression may compromise the effectiveness of subsequent immunotherapies, which are increasingly part of standard treatment regimens.

Remarkably, ultra-high dose rate (UHDR) “FLASH” RT—delivering the full radiation dose under 200 milliseconds at mean dose rates exceeding 40 Gy/s—has recently demonstrated the ability to selectively spare normal tissues while preserving tumor control in multiple preclinical models and its feasibility in humans has been tested in various clinical trials in the United States and in Europe(11–15). By minimizing the damage to healthy tissue while maintaining tumor efficacy, FLASH RT has been shown to significantly enhance the therapeutic ratio of RT and holds transformative potential for improving cancer treatment outcomes if successfully translated to clinical practice(16). To date, the exact mechanism for the FLASH effect remains unclear but a few hypotheses have been posited. One proposal is linked to oxygen dynamics(17). At physiologic oxygen levels, the intense burst of near-instantaneous ionization during FLASH RT rapidly depletes available oxygen, limiting the formation of reactive oxygen species (ROS) and thereby reducing oxidative DNA damage in normal cells(18–21). In contrast, CONV RT delivered over minutes allows time for tissue reoxygenation and sustained ROS generation, leading to greater normal tissue injury(22,23). This is likely caused by the differences in the radiation beam parameters from FLASH and CONV irradiation(16), where it has been reported that certain beam parameters are known to induce and optimize the FLASH effect(24). In FLASH RT, the entire radiation dose is delivered in a few pulses within a fraction of a second whereas in CONV RT the entire radiation dose is divided over hundreds to thousands of pulses in the span of minutes(25). In FLASH RT, the dose delivered per pulse, the instantaneous dose rate, and the mean dose rate are several orders of magnitude higher than those in CONV RT emphasizing the dramatic differences in the beam parameters that are employed between FLASH and CONV RT(26–28).

However, the oxygen depletion hypothesis does not account for the entirety of the FLASH effect due to studies observing UHDR RT being insufficient to completely deplete the oxygen in the treated volume(29–31), likely due to their dependence on a combination of the aforementioned radiation beam parameters, reoxygenation, and local tissue vasculature(32). A concomitant hypothesis for the FLASH effect is the substantial difference in the total blood volume irradiated under CONV versus FLASH RT(33,34). Under CONV RT, treatment times take minutes to deliver the total dose to the treatment volume allowing for a greater population of circulating lymphocytes to flow into the irradiated field and for a greater volume of blood lymphocytes to be irradiated, damaged, or killed, with the irradiated lymphocytes migrating to areas of the body outside of the radiation field. FLASH RT, however, has a total treatment time in under a fraction of a second thereby leaving a substantially smaller volume of blood irradiated due to having treatment times thousands of times shorter than conventional treatment times, thereby reducing inflammatory and fibrotic response in normal tissue(35,36). In fact, a study published by Jin et al. utilized a computational model to simulate dose rate effects on the depletion of circulating immune cells and has shown that a single fraction of 30 Gy depletes 90-100% of the circulating lymphocytes under CONV RT while 5-10% are depleted under FLASH RT indicating the therapeutic potential of FLASH in minimizing the irradiation of blood lymphocytes(33).

Due to the hypothesis that the total irradiated blood volume is linked to RIL, we examined the therapeutic potential of FLASH RT compared to CONV RT in the context of RIL for thoracic irradiations, both single- and multi-fraction deliveries, along with the corresponding mechanism behind differences in their effects.

## Materials and Methods

### Mice

All animal procedures were conducted according to the Guide for the Care and Use of Laboratory Animals, prepared by the Institute of Laboratory Animal Resources, National Research Council and National Academy of Sciences, and the MD Anderson Cancer Center Institutional Animal Care and Use Committee. The experimental protocols were approved by the Institutional Animal Care and Use Committee (IACUC) at The University of Texas MD Anderson Cancer Center. C57BL/6 female mice at 8 weeks old were purchased from Jackson Laboratory (Bar Harbor, Maine, USA). Mice were acclimatized for a minimum of 3 days prior to performing an experiment. The mice had access to standard chow (PicoLab Rodent Diet 20, no. 5053; PMI Nutrition International, St. Louis, MO, United States) and acidified water ad libitum. Five mice were housed per ventilated cage in a 12/12-hour light/dark cycle.

### Irradiation Setup and Treatment

All irradiations were performed at MD Anderson Cancer Center on the IntraOp Mobetron electron linear accelerator (IntraOp Medical, Sunnyvale, CA, USA) that can deliver 9-MeV electrons at CONV and UHDR settings(37–41). A custom-made mouse cradle was designed and 3D-printed to immobilize mice while they were sedated with isoflurane in ambient air (3.5% for anesthetic induction and 1.5% for maintenance, with total anesthesia time < 5 minutes).

Mouse positioning in the cradle was verified by CT imaging on a small animal radiation research platform (SARRP, Xstrahl Inc., Swanee, GA, USA) as mentioned in previous *in vivo studies* conducted on the same Mobetron(26,42). A custom-made collimator was produced and seated at the exit window of the Mobetron, as described elsewhere(42), to allow for sedation and reproducible setup placement for full and consistent total thoracic irradiation to a field size of 4 x 2 cm^2^. The irradiation setup was calibrated for each irradiation condition listed in Table S1 to a depth of 8 mm in solid water.

Dosimetric calibration was performed using the protocols mentioned in previous studies with dose-rate-independent Gafchromic EBT3 film(43) paired with beam current transformers(44), the latter allowing for real-time measurement of the individual radiation beam parameters for each delivery. During the UHDR irradiations, the beam current transformers were used to monitor the dose delivery and to log all radiation beam parameters on a per-pulse basis for each individual mouse. All CONV irradiations were performed at a set mean dose rate of 0.3 Gy/s and a dose per pulse of 10 mGy and were monitored using the internal ion chambers inside the Mobetron.

To evaluate the effects of FLASH RT on lymphocyte preservation compared to CONV RT, we utilized the Mobetron to deliver thoracic RT to C57BL/6 mice. Eight-week-old mice received a single fraction 17 Gy dose of either CONV or FLASH RT to the whole thorax with untreated mice serving as controls. To better replicate clinical radiation treatment conditions, a fractionated low-dose radiation regimen was employed. Eight-week-old C57BL/6 mice received thoracic irradiation with either CONV or FLASH RT with 2 Gy delivered per day for five consecutive days (total 10 Gy; 4 × 2 cm^2^ field). SHAM-treated mice served as controls. Detailed dose rates and beam parameters are provided in Supplementary Table 1.

### Flow Cytometry Analysis

Peripheral blood from mice was collected at the longitudinal time points indicated in Figure 1 via the facial vein using lithium-heparin (LH)-coated microvettes (Sarstedt, Nümbrecht, Germany) for both single-fraction and multi-fraction treatments. Samples were incubated on ice for 45 minutes with a cocktail of fluorophore-conjugated primary antibodies. Red blood cells were then lysed using RBC lysis buffer (BioLegend) for 15 minutes. Following two washes with phosphate-buffered saline (PBS), cells were fixed with IC Fixation Buffer (Invitrogen) and transferred to round-bottom polystyrene tubes with cell-strainer caps (Falcon). CountBright™ Absolute Counting Beads (Invitrogen) were added to each sample prior to acquisition on a BD LSRFortessa™ Cell Analyzer (BD Biosciences). Data were analyzed using FlowJo software.

**Figure 1.**
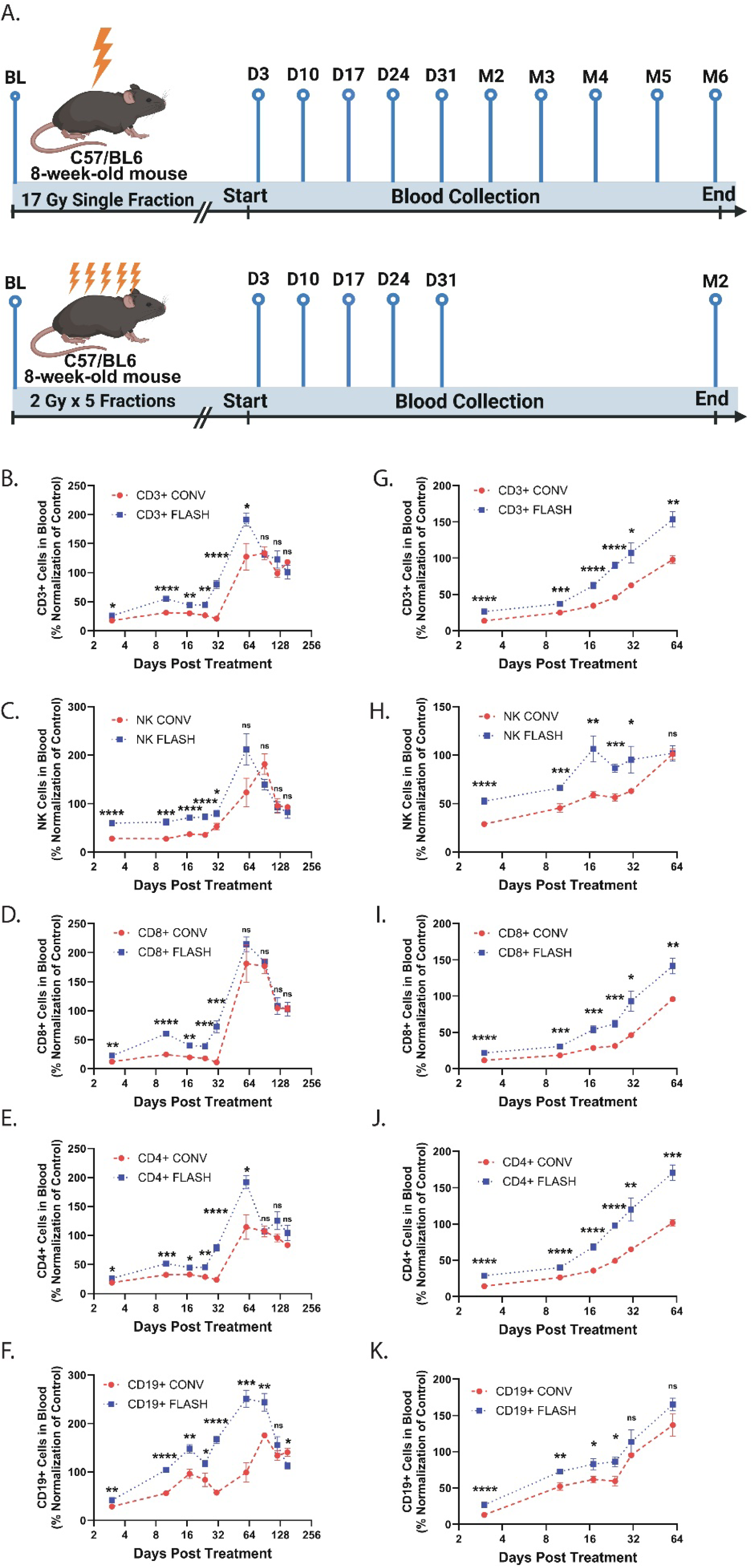
Thoracic FLASH Radiotherapy Preserves Peripheral Lymphocytes and Enhances Lymphopenia Recovery in Mice. (A) Schematic of experimental design for single- and multi-fraction thoracic irradiation with blood collected at the indicated timepoints. (B–F) Eight-week-old C57BL/6 mice received a single 17 Gy dose of conventional (CONV) or FLASH RT to the whole thorax (4 × 2 cm^2^). Non-irradiated mice served as controls. (G–K) A separate cohort received a multi-fraction regimen of 2 Gy/day for 5 consecutive days of CONV or FLASH RT to the thorax, with controls included. Peripheral blood was collected via facial vein at the indicated time points and analyzed by flow cytometry to quantify lymphocyte populations (CD3+, NK, CD8+, CD4+, and CD19+). Sample sizes: Single-fraction group—N = 8 mice/group (baseline to day 24); N = 5 mice/group (day 31 to month 5). Multi-fraction group—N = 7 mice/group (baseline to day 17); N = 5 mice/group (day 24 to month 2). Data are shown as mean ± standard deviation. Statistical significance: *p < 0.05, **p < 0.01, ***p < 0.001, ****p < 0.0001. Data are representative of at least three independent experiments. Abbreviations: BL, baseline; D, day; M, month; CTR, control; CONV, conventional; ns, not significant.

### Masson’s Trichrome Staining Assay

Masson’s Trichrome staining was performed using the Trichrome Stain Kit (Abcam, ab150686) following the manufacturer’s protocol. Briefly, mouse lung and heart were harvested and fixed in 10% buffered formalin (Fisherbrand) 6-months post-irradiation. Paraffin embedding and hematoxylin and eosin (H&E) staining were conducted by the MD Anderson Research Histology Core Laboratory.

Prior to staining, all reagents and materials were equilibrated to room temperature with gentle agitation. Tissue sections were deparaffinized and rehydrated in distilled water. Bouin’s fluid was preheated to 60°C in a fume hood, and slides were incubated in the preheated Bouin’s solution for 60 minutes, followed by a 10-minute cooling period. Sections were then rinsed thoroughly in running tap water until clear and rinsed once in distilled water. Equal volumes of Weigert’s Iron Hematoxylin Solutions A and B were mixed to prepare the working solution. Slides were stained with the working hematoxylin for 5 minutes and rinsed in running tap water for 2 minutes. Next, Biebrich Scarlet/Acid Fuchsin solution was applied for 15 minutes, followed by a rinse in distilled water. Slides were differentiated in Phosphomolybdic/Phosphotungstic Acid solution for 15 minutes, then directly stained with Aniline Blue solution for 10 minutes without intermediate rinsing. After another rinse in distilled water, slides were treated with 1% Acetic Acid for 5 minutes.

Sections were dehydrated rapidly through two changes of 95% ethanol, followed by two changes of absolute ethanol, cleared in xylene, and mounted using VectaMount Mounting Medium (Vector Laboratories, H-5000). Stained sections were imaged using a Leica Aperio CS2 Scanscope (Leica Biosystems) and analyzed with ImageScope software (Leica Biosystems).

### Immunofluorescence Staining

Frozen spleen tissue sections from mice two-months post-treatment, along with hematoxylin and eosin (H&E) staining, were prepared by the MD Anderson Research Histology Core Laboratory. Fixation and permeabilization were performed by immersing the frozen sections in cold acetone for 15 minutes, as previously described(45). Following fixation, slides were rinsed with phosphate-buffered saline (PBS) and blocked for 1 hour at room temperature using 4% fish gelatin (Biotium) diluted in PBS. Tissues were then incubated overnight at 4°C with the following primary antibodies, each diluted in 4% fish gelatin: rat anti-CD45 (BD Pharmingen), rabbit anti-phospho-Chk1 (Ser345) (Cell Signaling Technology), mouse anti-phospho-Stat3 (Tyr705) (B-7) (Santa Cruz Biotechnology), goat anti-PD-1 (R&D Systems), and goat anti-PD-L1/B7-H1 (R&D Systems).

After PBS washes, sections were incubated for 2 hours at RT with the following secondary antibodies, all diluted in 4% fish gelatin: donkey anti-rat Alexa Fluor 488, anti-rabbit Alexa Fluor 594, anti-mouse Alexa Fluor 555, and anti-goat Alexa Fluor 750 (all from Abcam). Slides were then washed with PBS and mounted using Fluoro-Gel II with DAPI (Electron Microscopy Sciences). Images were acquired using a Vectra Polaris Quantitative Slide Scanner (PerkinElmer) and analyzed with Phenochart software version 2.2.0 (Akoya Biosciences).

### Western Blot Analysis

Treated mouse spleen tissues were collected 2-months post-irradiation, and protein extracts were prepared for Western blotting using protocols described previously(45–47). Briefly, tissues were lysed in Cell Lysis Buffer (Cell Signaling Technology, Danvers, MA) supplemented with protease and phosphatase inhibitor cocktails (Thermo Scientific) and incubated on ice for 30 minutes. Lysates were then sonicated and clarified by centrifugation at 13,000 × g for 10 minutes at 4°C. The resulting supernatants were collected and denatured by boiling in SDS loading buffer (Cell Signaling Technology).

Proteins were separated by electrophoresis on 10% SDS-polyacrylamide gels (Bio-Rad Laboratories, Hercules, CA), then transferred to nitrocellulose membranes (Whatman Schleicher & Schuell, Keene, NH). Membranes were blocked and incubated overnight at 4°C with the following primary antibodies: mouse anti-Chk1, rabbit anti-phospho-Chk1 (Ser345), mouse anti-Stat3, rabbit anti-phospho-Stat3 (Tyr705), rabbit anti-α-Tubulin (all from Cell Signaling Technology), and goat anti-PD-1 and goat anti-PD-L1/B7-H1 (both from R&D Systems).

Following washes, membranes were incubated for 2 hours at room temperature with the appropriate horseradish peroxidase (HRP)-conjugated secondary antibodies: anti-rabbit, anti-mouse, or anti-goat (Thermo Scientific). Signal detection was performed using the SuperSignal West Dura Extended Duration Substrate (Thermo Scientific), and protein bands were visualized using either a Kodak Medical X-Ray Processor 104 (Eastman Kodak, Rochester, NY) or a ChemiDoc Touch Imaging System (Bio-Rad). Images were processed and analyzed using Adobe Photoshop.

### Statistical Analysis

Statistical analyses were performed using GraphPad Prism software. Comparisons between two groups were conducted using a two-tailed unpaired t-test. Data are presented as mean ± standard deviation (SD), and p-values less than 0.05 were considered statistically significant.

## Results

### Single- and Multi-Fraction Thoracic FLASH Radiation Therapy (RT) Spares Lymphocytes, Mitigates Fibrosis, and Promotes Recovery from Radiation-Induced Lymphopenia in Mouse Peripheral Blood

Longitudinal blood sampling was performed at the indicated time points (Figure 1A), and lymphocyte subsets were analyzed via flow cytometry. As shown in Figures 1B–1F, single-fraction CONV RT induced severe and sustained lymphopenia across all major lymphocyte populations—including CD3^⁺^, CD4^⁺^, CD8^⁺^ T cells, NK cells, and CD19^⁺^ B cells—with minimal recovery over time, aside from a modest resurgence of CD19^⁺^ B cells by two months post-treatment. In contrast, single-fraction FLASH RT also triggered an initial decline in lymphocyte counts by day 3; however, a progressive recovery was observed, reaching near-baseline levels by day 31 (nearly a month faster recovery than the single-fraction CONV-treated group). Notably, lymphocyte counts in single-fraction FLASH-treated mice were significantly higher (p < 0.05, 1.2-6.5x higher) than those in the single-fraction CONV group across all subsets within the first 2 months post-irradiation, highlighting FLASH RT’s capacity to mitigate acute lymphopenia and accelerate immune reconstitution post-irradiation. At six-months post-irradiation, lungs and hearts from the single-fraction treated mice were collected and stained with Masson’s Trichrome to assess for late radiation-induced tissue damage. As shown in Figure S1, single-fraction CONV RT led to pronounced fibrosis in both organs, whereas single-fraction FLASH RT significantly reduced fibrosis (4-5x higher relative collagen production in CONV-treated mice compared to FLASH-treated mice, p < 0.0001), indicating attenuated long-term tissue injury in mice treated with FLASH RT.

Figures 1G–1K show similar effect in the context of fractionated treatment. Fractionated CONV RT induced sustained lymphopenia in all major lymphocyte subsets, with limited recovery over a two-month period post-irradiation, mirroring the single-fraction CONV RT outcomes. In contrast, fractionated FLASH RT caused an initial drop in lymphocyte counts followed by steady recovery, with levels returning to near-control values by day 24— approximately one week earlier than recovery observed in the single-fraction FLASH RT group. Furthermore, lymphocyte levels in the fractionated FLASH group were found to be significantly higher (p < 0.05, 1.01-2.04x higher) than those in the fractionated CONV group at multiple time points evaluated with the amount of time for the lymphocyte counts to reach to the level near baseline controls being nearly a month faster in the FLASH-treated group compared the CONV-treated group. Together, these findings demonstrate that both single-fraction and multi-fraction FLASH RT significantly mitigate RIL, radiation-induced fibrosis, and promote a more rapid and robust recovery of lymphocytes compared to CONV RT.

### Single- and Multi-Fraction Thoracic FLASH RT Significantly Reduces Radiation-Induced Lymphocyte Apoptosis

To explore the mechanism underlying the lymphocyte-sparing effect of FLASH RT, we assessed RIL apoptosis in peripheral blood, given the well-established role of ionizing radiation in triggering apoptotic cell death(48). Peripheral blood was collected at 16 hours, 36 hours, 3 days, and 10 days post-irradiation via facial vein bleed. Apoptotic lymphocytes were identified using Annexin V staining in conjunction with surface markers for CD45+, CD3+, CD4+, CD8+, NK cells, and CD19^⁺^ B cells, and were quantified by flow cytometry.

As shown in Figures 2A–2F, single-fraction CONV RT resulted in a significant increase in apoptosis across all lymphocyte subsets as early as 16 hours post-irradiation, with peak apoptotic cell levels observed at 36 hours and 3 days post-irradiation. Meanwhile single-fraction FLASH RT also induced an increase in apoptotic lymphocytes compared to controls, the levels of apoptotic CD3^⁺^, CD4^⁺^, CD8^⁺^ T cells, NK cells, and B cells were significantly lower in FLASH-treated mice than in CONV-treated mice at each of the early time points up to 3 days post-irradiation (p < 0.05, 1.5-6.1x lower in FLASH RT compared to CONV RT). By the 10^th^ day post-irradiation, the levels of apoptotic cells returned to baseline comparable to SHAM in both treatment groups. These results indicate that FLASH RT reduces the magnitude and severity of acute RIL apoptosis, potentially contributing to the preservation of circulating lymphocytes after irradiation.

**Figure 2.**
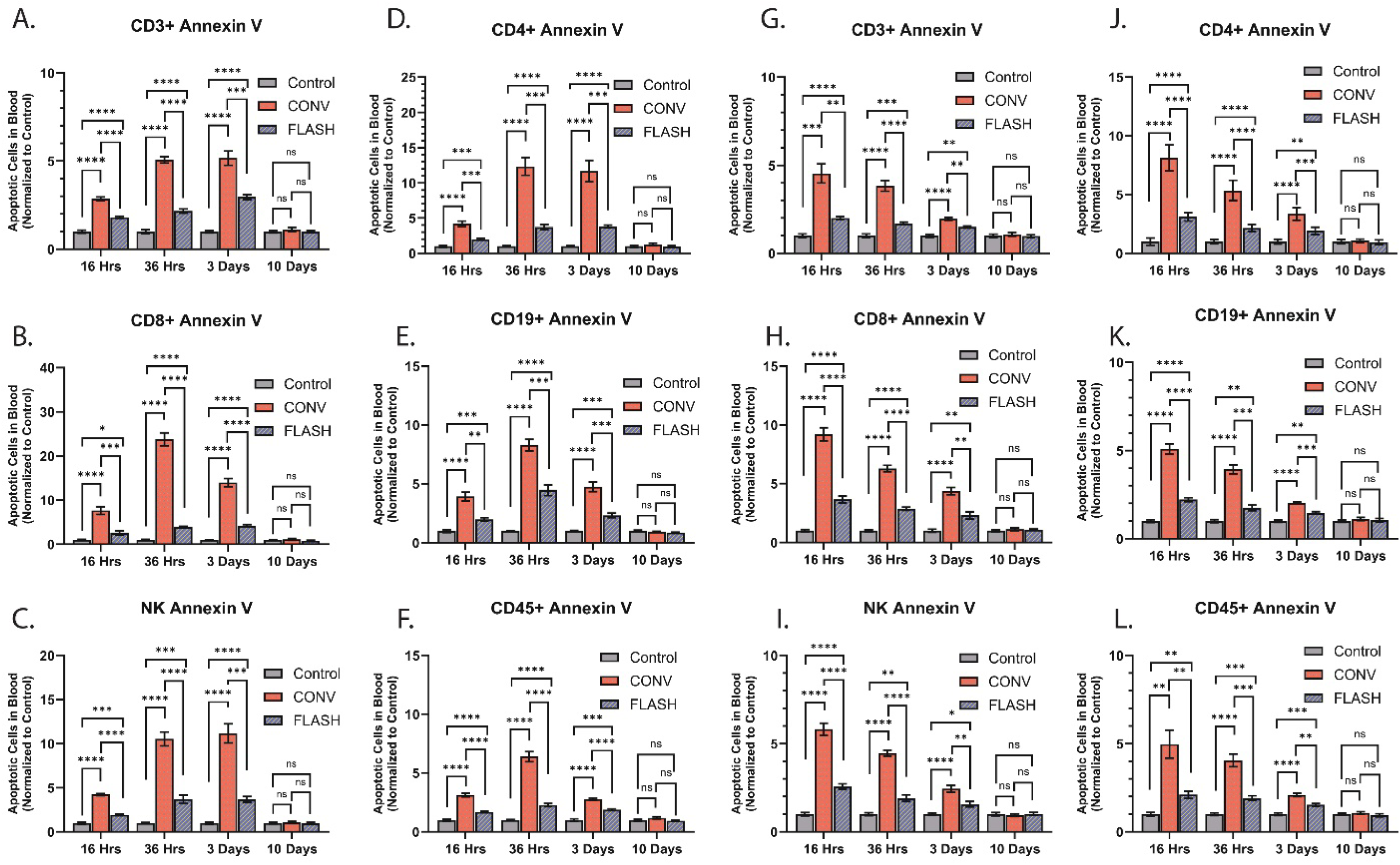
Single- and Multi-Fraction Thoracic FLASH Radiotherapy Significantly Reduces Radiation-Induced Lymphocyte Apoptosis. (A–F) Eight-week-old C57BL/6 mice received a single 17 Gy dose of conventional (CONV) or FLASH radiotherapy (RT) to the whole thorax. (G–L) A separate cohort received fractionated RT (2 Gy/day for 5 consecutive days) using CONV or FLASH RT; non-irradiated mice served as controls. Peripheral blood was collected via facial vein at 16 hours, 36 hours, 3 days, and 10 days post-irradiation. Apoptosis in lymphocyte subpopulations (CD45^⁺^, CD3^⁺^, CD4^⁺^, CD8^⁺^, NK, and B cells) was assessed by Annexin V staining and analyzed via flow cytometry. Sample size: N = 5 mice per group. Data are presented as mean ± standard deviation. Statistical significance: *p < 0.05, **p < 0.01, ***p < 0.001, ****p < 0.0001.

To determine whether these protective effects also extend to fractionated FLASH RT, we evaluated apoptosis following multi-fraction treatment. As shown in Figures 2G–2L, fractionated CONV RT similarly induced a marked increase in apoptotic cells across all lymphocyte subsets, with peak levels occurring 16 hours following the final dose fraction. Although fractionated FLASH RT also led to an increase in the number of apoptotic cells relative to the control group, the population of apoptotic CD3^⁺^, CD4^⁺^, CD8^⁺^ T cells, NK cells, and B cells was consistently and significantly lower compared to those in the fractionated CONV RT group at all early time points up to 3 days post-irradiation (p < 0.05, 1.3-2.6x lower in FLASH RT compared to CONV RT).

These findings provide a potential mechanistic insight into the higher lymphocyte counts and accelerated recovery from lymphopenia observed in FLASH-treated mice. Collectively, the data suggest that the lymphocyte-sparing effect of FLASH RT is at least partially mediated by a reduction in radiation-induced apoptosis during the acute post-irradiation period for both single-fraction and multi-fraction irradiation regimens.

### Conventional, but Not FLASH, Thoracic Radiotherapy Induces Sustained Immunosuppression via PD-1 and PD-L1 Expression on Lymphocytes Following Lymphopenia Recovery

RT induces immune suppression through both direct depletion of circulating lymphocytes and through modulating the microenvironment that generates chronic inflammation and upregulation of immune suppressor cells such as T-regulatory cells (Tregs) (49,50). We evaluated a marker of chronic inflammation that leads to immune suppression is induction of immune checkpoint proteins programmed-death-1 (PD-1) and programmed-death-ligand-1 (PD-L1). As shown in Fig. 3, both CONV and FLASH thoracic RT led to a marked reduction in PD-1^⁺^ and PD-L1^⁺^ lymphocyte numbers 3 days post-irradiation, consistent with acute lymphopenia. By day 24, lymphocyte levels recovered to near-baseline values in both single- and multi-fraction treatment groups for both CONV and FLASH RT (with exception to CD19+ cells for the CONV treated group).

**Figure 3.**
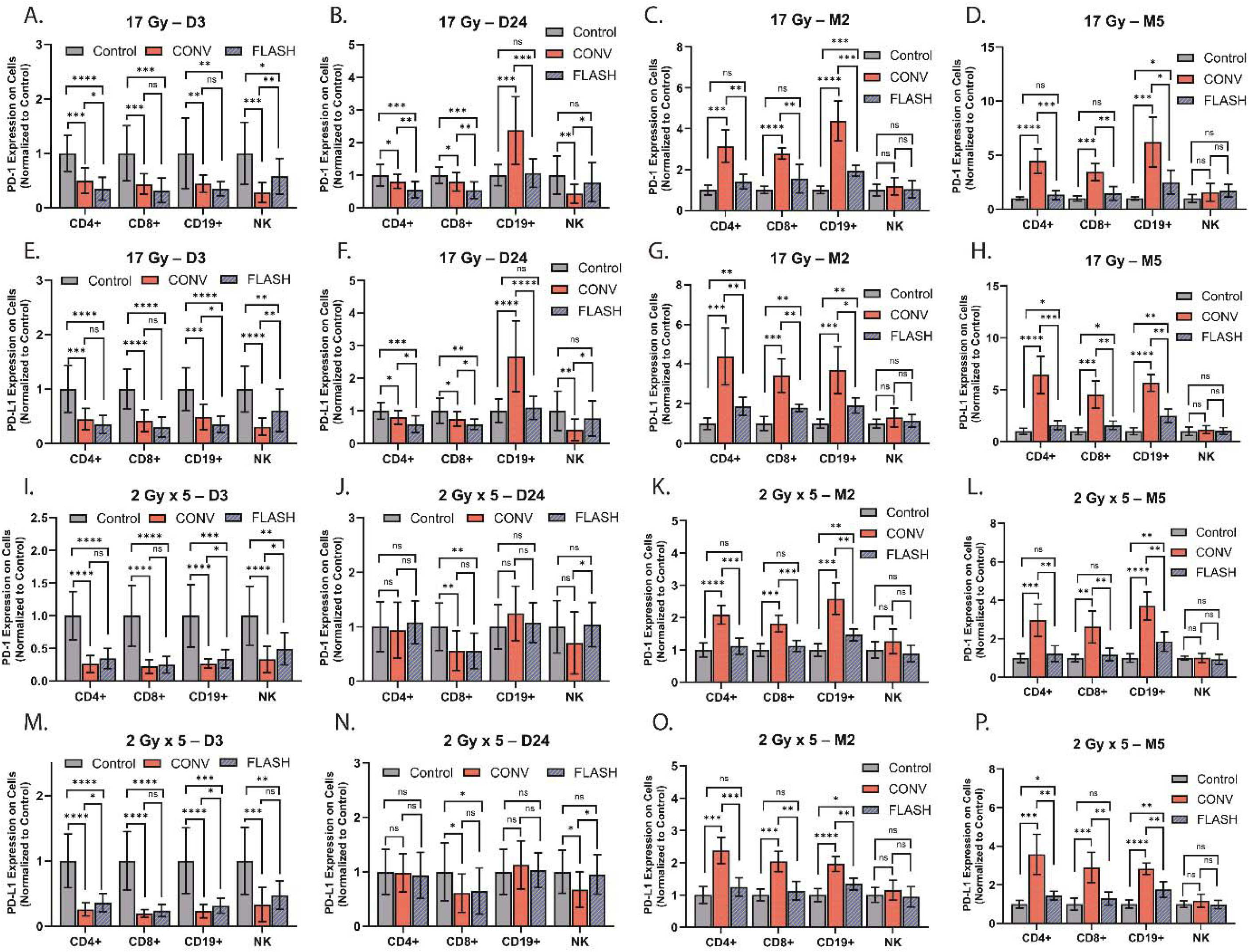
Conventional, but Not FLASH, Thoracic Radiotherapy Induces Sustained PD-1 and PD-L1 Expression on Lymphocytes Following Lymphopenia Recovery. Eight-week-old C57BL/6 mice received either a (A–H) single-fraction 17 Gy dose or (I–P) multi-fraction (2 Gy x 5) thoracic RT using conventional (CONV) or FLASH RT; non-irradiated mice served as controls. Peripheral blood was collected via facial vein at day 3, day 24, month 2, and month 5 post-irradiation, as indicated. PD-1 and PD-L1 expression was assessed by flow cytometry on CD4^⁺^, CD8^⁺^, CD19^⁺^, and NK cell populations. Sample size: N = 5 mice per group. Data are presented as mean ± standard deviation. Statistical significance: *p < 0.05, **p < 0.01, ***p < 0.001, ****p < 0.0001.

However, long-term expression patterns differed significantly. As illustrated in Fig.3A-3H a single 17 Gy fraction of CONV RT caused a persistent upregulation of PD-1 and PD-L1 expression in CD4^⁺^ and CD8^⁺^ T cells, as well as CD19^⁺^ B cells, at both 2- and 5-months post-treatment (p < 0.05, 1.9-4.1x higher in CONV RT relative to FLASH RT). Notably, NK cells showed no significant changes in PD-1 or PD-L1 expression. In contrast, single-fraction FLASH RT had a significantly smaller impact on PD-1 and PD-L1 expression across these lymphocyte subsets at 2- and 5-months post-irradiation compared to CONV RT.

A similar trend was observed in the multi-fraction setting (2 Gy × 5 days; Figures 3I-3P), where fractionated CONV RT induced sustained PD-1 and PD-L1 expression on T and B cells (p < 0.01, 1.5-2.5x higher in CONV RT relative to FLASH RT), but not on NK cells. FLASH RT under the same fractionated regimen produced only a modest increase in PD-L1 expression on B cells, with little to no effect on T or NK cells. These findings suggest that CONV RT—both single-fraction and multi-fraction regimen—induces prolonged PD-1 and PD-L1 expression on lymphocytes, which may contribute to a lasting immunosuppressive state several months post-irradiation for both single-fraction and multi-fraction treatment regimens.

### Conventional, but Not FLASH, Thoracic RT Induces a Sustained Increase in Treg Cells Following Lymphopenia Recovery

As shown in Figures 4A–4D, both single-fraction 17 Gy doses of CONV and FLASH RT resulted in a significant early decline in Treg cell populations at day 3 and day 24 post-irradiation. However, by 2- and 5-months post-irradiation, mice treated with single-fraction CONV RT exhibited a significantly elevated increase in Treg cells compared to the control group (p < 0.001). In contrast, Treg cell levels in the single-fraction FLASH RT group showed some elevation but were significantly less than the single-fraction CONV RT group (p < 0.01, 2.3-2.9x lower than CONV RT) and remained comparable to the levels found in control mice, indicating the absence of a delayed Treg rebound following FLASH treatment.

**Figure 4.**
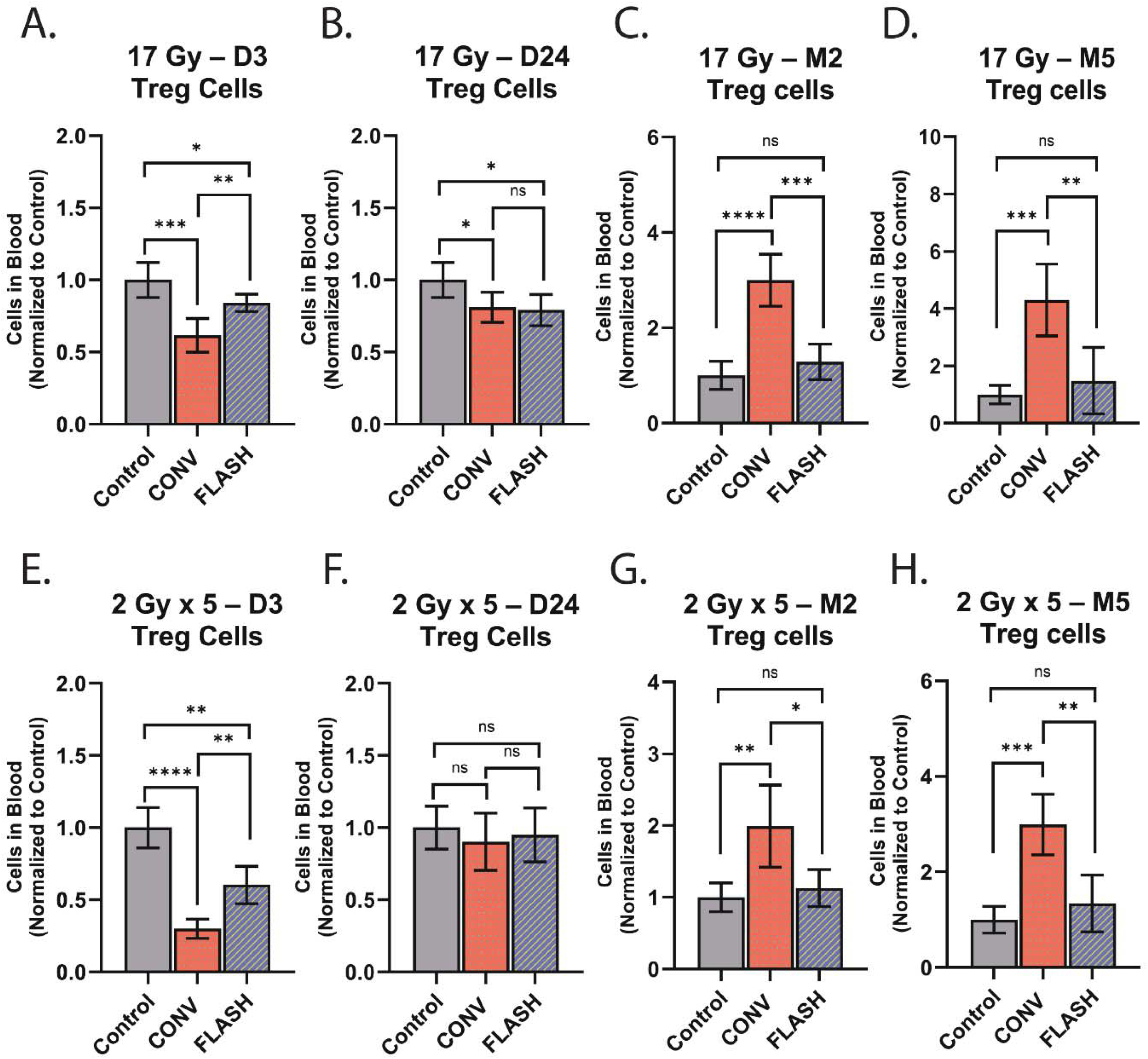
Conventional, but Not FLASH, Thoracic Radiotherapy Increases Regulatory T Cell Populations Following Lymphopenia Recovery. Eight-week-old C57BL/6 mice were subjected to either (A–D) single-fraction 17 Gy or multi-fraction (E–H) 2 Gy x 5 conventional (CONV) or FLASH radiotherapy to the whole thorax. Non-irradiated mice served as controls. Peripheral blood was collected via facial vein at day 3, day 24, month 2, and month 5 post-irradiation, as indicated. Regulatory T cells (Tregs) were quantified by flow cytometry. Sample size: N = 5 mice per group. Data are presented as mean ± standard deviation. Statistical significance: *p < 0.05, **p < 0.01, ***p < 0.001, ****p < 0.0001.

In the fractionated RT setting, both CONV and FLASH groups showed a similar early decline in Treg cell counts at day 3, followed by recovery to baseline levels by day 24 post-irradiations (Fig. 4E-4H). Notably, at 2- and 5-months post-irradiation, fractionated CONV RT again led to a sustained elevation in Treg cells relative to the control group (p < 0.01), mirroring the pattern observed in the single-fraction cohort. However, mice treated with fractionated FLASH RT maintained Treg cell levels that were significantly lower than those in the CONV group (p < 0.05, 1.8-2.2x lower than CONV RT) and were nonsignificant from the control group (p > 0.05).

These findings suggest that CONV RT—both single and fractionated treatment regimens—leads to a long-term increase in immunosuppressive Treg cells, even after apparent recovery from RIL. In contrast, FLASH RT prevents this delayed immunosuppressive effect, preserving an immune profile comparable to the control cohort several months after treatment.

### FLASH RT Attenuates Radiation-Induced Immunosuppression via Inhibition of the Chk1– STAT3 Pathway

To investigate the molecular mechanism underlying the differential immunosuppressive effects of CONV versus FLASH RT, we explored the signaling axis responsible for radiation-induced PD-1 and PD-L1 expression in lymphocytes. While the regulatory mechanisms remain largely undefined, STAT3 is a known transcription factor for both Pdcd1 (PD-1) and Cd274 (PD-L1) genes(51–53) .

We first examined whether STAT3 activation was associated with increased PD-1 and PD-L1 expression in splenic lymphocytes post-radiation. As shown in Figures 5A and 5B, the population of CD45^⁺^PD-1^⁺^p-STAT3^⁺^ and CD45^⁺^PD-L1^⁺^p-STAT3^⁺^ triple-positive cells were significantly higher (p < 0.01, 1.7-2.5x higher) in CONV-treated mice compared to both FLASH-treated and control groups in both the single- and multi-fraction treatment, indicating that PD-1 and PD-L1 upregulation is associated with radiation-induced STAT3 activation.

**Figure 5.**
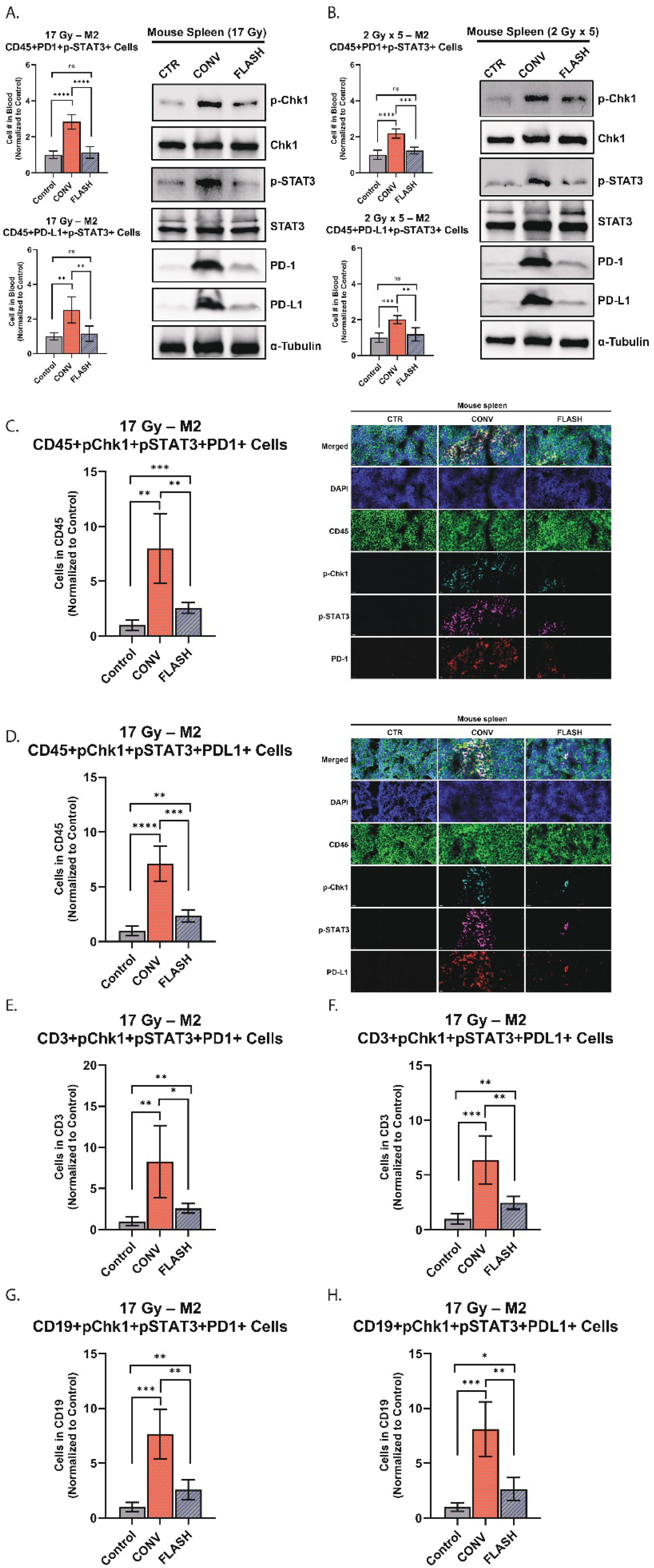
FLASH Radiotherapy Reduces Radiation-Induced Immunosuppression via the Chk1–STAT3 Signaling Axis. Eight-week-old C57BL/6 mice received either (A) single-fraction (17 Gy) or (B) multi-fraction 2 Gy x 5 thoracic irradiation using conventional (CONV) or FLASH radiation therapy (RT); non-irradiated mice served as controls. Peripheral blood was collected via facial vein at 2 months post-irradiation. Flow cytometry was used to quantify CD45^⁺^PD-1^⁺^p-STAT3^⁺^ and CD45^⁺^PD-L1^⁺^p-STAT3^⁺^ immune cells. Spleens from the same mice were harvested at month 2 for additional analyses. Western blot analysis was performed on spleen protein extracts to assess expression of pathway-related proteins of mice treated with single-fraction FLASH and CONV RT. (C–D) Immunofluorescence staining was performed on spleen sections of mice treated with single-fraction FLASH and CONV RT harvested 2 months post-irradiation; representative images show expression of indicated markers. Scale bar, 10 µm. (C-H) Flow cytometry was used to analyze the following cell populations from single-fraction treated mice: CD45^⁺^p-Chk1^⁺^p-STAT3^⁺^PD-1^⁺^, CD45^⁺^p-Chk1^⁺^p-STAT3^⁺^PD-L1^⁺^, CD3^⁺^p-Chk1^⁺^p-STAT3^⁺^PD-1^⁺^, CD3^⁺^p-Chk1^⁺^p-STAT3^⁺^PD-L1^⁺^, CD19^⁺^p-Chk1^⁺^p-STAT3^⁺^PD-1^⁺^, and CD19^⁺^p-Chk1^⁺^p-STAT3^⁺^PD-L1^⁺^. Sample size: N = 5 mice per group. Data are presented as mean ± standard deviation. Statistical significance: *p < 0.05, **p < 0.01, ***p < 0.001, ****p < 0.0001.

Since STAT3 can be phosphorylated downstream of DNA damage response pathways—particularly via Chk1, which is activated by radiation-induced DNA damage(54,55)—we hypothesized that CONV RT induces phosphorylation of Chk1, leading to downstream activation of STAT3 and transcriptional upregulation of PD-1 and PD-L1. To test this, we analyzed protein extracts from splenic tissues collected two months post-irradiation via western blot. As shown in Figures 5A and 5B, CONV RT (both single- and multi-fraction treatment) induced high levels of phosphorylated Chk1 (p-Chk1) and phosphorylated STAT3 (p-STAT3), with concomitantly elevated PD-1 and PD-L1 expression. In contrast, FLASH RT had greatly attenuated levels of p-Chk1, p-STAT3, and PD-1 and PD-L1 levels relative to CONV, and only marginally higher than the non-irradiated group.

To confirm that these signaling events occur within the same lymphocyte populations, we performed immunofluorescence staining of spleen sections collected two months post-treatment for the single-fraction treated groups. CONV RT demonstrated significantly higher PD-1 and PD-L1 expression in CD45^⁺^ lymphocytes (Figures 5C and 5D), CD3^⁺^ lymphocytes (Figures 8E, 8F, S2A, and S2B), and CD19^⁺^ lymphocytes (Figures 8G, 8H, S2C, and S2D). Importantly, the majority of PD-1^⁺^ and PD-L1^⁺^ cells also co-expressed p-Chk1 and p-STAT3 for CD45^⁺^, CD3^⁺^, and CD19^⁺^ lymphocytes, a pattern that is much reduced in the FLASH-treated mice.

Together, these results support a model in which CONV RT induces chronic inflammation and immunosuppressive signaling in lymphocytes through the Chk1–STAT3 axis, promoting PD-1 and PD-L1 expression. In contrast, FLASH RT mitigates this pathway, thereby preventing long-term radiation-induced immunosuppression.

## Discussion

Our findings demonstrate that thoracic FLASH RT, delivered as either a single-fraction 17 Gy dose or a multi-fraction 2 Gy × 5 day regimen, provides significant immunological advantages over CONV RT of identical regimen in a preclinical mouse model. FLASH RT not only minimized radiation-induced fibrosis and lymphopenia, even several months post-treatment but also enabled a faster and more complete recovery of lymphocyte populations, including CD3^⁺^, CD4^⁺^, CD8^⁺^, NK, and CD19^⁺^ B cells. These effects were observed in the acute time frame—as early as 3 days post-treatment—and persisted longitudinally, with lymphocyte counts returning to near-baseline levels by day 31 in the single-fraction FLASH group and by day 24 in the fractionated FLASH group indicating that the FLASH effect is maintained even when using a standard-of-care fractionation regime(11), which is rarely reported in the literature except for one previous study documenting the FLASH effect in whole brain irradiation(56). In contrast, CONV RT resulted in sustained lymphopenia, with little to no recovery in T or NK cell compartments and delayed rebound of B cells, consistent with previously reported immunosuppressive effects of standard radiotherapy regimens(5,57–59).

The protective effects of FLASH RT were further substantiated by reduced lymphocyte apoptosis at acute time points following irradiation. Flow cytometric analysis revealed significantly lower Annexin V staining across all major lymphocyte subsets in FLASH-treated mice compared to those receiving CONV RT, suggesting that FLASH RT may limit apoptotic cell death via mechanisms distinct from CONV RT damage responses. Previous studies investigating the therapeutic efficacy of FLASH have reported that FLASH RT results in less apoptosis and senescence in normal tissue compared to CONV RT(35,60–62), produces fewer dicentric chromosomes in human blood lymphocytes(63), induces significantly lower levels of DNA damage in peripheral blood lymphocytes(64) and generates fewer clustered DNA damage sites compared to CONV irradiation in vitro(65) indicating a potential link for the significantly lower levels of apoptosis detected in mice treated with FLASH RT compared to CONV RT.

Importantly, the long-term immunological landscape was markedly different between treatment modalities. Both single-fraction and multi-fraction CONV RT induced sustained increases in the Treg population and enhanced expression of the immune checkpoint proteins PD-1 and PD-L1 on CD4^⁺^, CD8^⁺^, and B cells—changes consistent with a systemic immunosuppressive phenotype(66–70). These immunological alterations were evident during the apparent recovery of total lymphocyte counts at several months post-irradiation. In contrast, FLASH RT did not lead to such changes; the Treg levels remained similar to those in the untreated controls at the later timepoints (2- and 5-months post-irradiation) and the PD-1 and PD-L1 expression remained significantly lower than the CONV RT mice at all time points examined (with exception to the NK cells at 2- and 5-months post-irradiation).

Mechanistically, our data is the first to implicate the Chk1-STAT3 signaling axis in the differential outcomes observed. We found that CONV RT activated Chk1 phosphorylation and subsequently induced STAT3 phosphorylation, both in splenic tissue and within PD-1^⁺^ or PD-L1^⁺^ lymphocyte populations. This pathway is known to regulate transcription of PD-1 and PD-L1 and likely mediates the observed checkpoint upregulation following CONV RT(71–74). FLASH RT, by contrast, did not activate this signaling axis, providing a potential molecular basis for its immune-sparing properties. These findings build upon existing literature suggesting that FLASH RT may preserve normal tissue integrity through a variety of mechanisms potentially involving redox modulation, transient oxygen depletion, or altered DNA damage signaling(20,63,64,75–78).

Our study is limited by the use of only nontumor-bearing animals. Tumor-bearing animals inherently have limited survival which would preclude any meaningful assessment of long-term lymphocyte recovery or sustained immune suppression. Nevertheless, we believe our findings establish the foundation for future studies to evaluate the long-term effects of both treatment modalities in tumor-bearing animals. Although we observed a strong correlation between chronic immune signaling in lymphocytes and PD-1/PD-L1 expression in CONV-treated mice, causality cannot be inferred. One plausible explanation is that radiation-damaged lymphocytes circulating through the thorax, which evade apoptosis, persist and later manifest as chronically inflamed cells within the spleen. Additional preclinical track-tracing studies will be needed to determine if this is indeed the case. Finally, the precise mechanisms of how dose rate can influence chronic inflammatory signaling in cells outside of the radiation field needs to be further elucidated in future studies.

## Conclusion

Taken together, these results highlight the potential of FLASH RT to overcome one of the major negative impacts of RT—radiation-induced lymphopenia and immunosuppression. By minimizing lymphocyte apoptosis, preserving immune cell populations, and preventing chronic checkpoint activation that induces immunosuppression, FLASH RT may better preserve immune healthy in the context of cancer therapy, particularly when RT is combined with immunotherapies, where lymphocyte function is critical for treatment efficacy. Future studies should explore the translational potential of FLASH RT in clinical settings, including its effects on tumor-infiltrating lymphocytes, response to immunomodulatory agents, and long-term safety(17). Additionally, further mechanistic work is warranted to evaluate the temporal dynamics and molecular mediators of FLASH-specific immune preservation.

---

**Figure S1.**
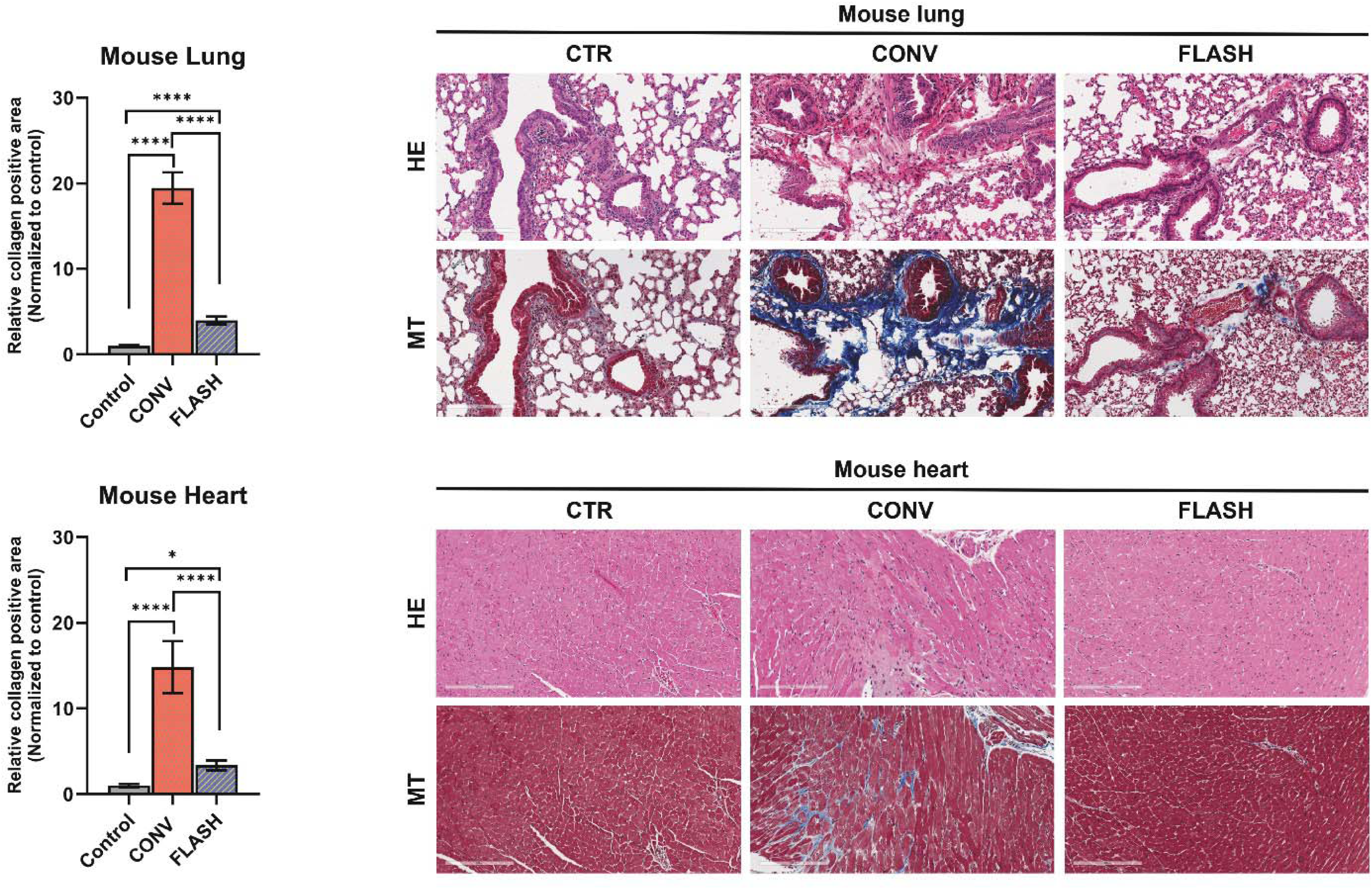
FLASH Radiotherapy reduces lung and heart fibrosis in mice. Eight-week-old C57BL/6 mice received single-fraction thoracic irradiation (17 Gy) using either conventional (CONV) or ultra-high dose rate FLASH radiotherapy; non-irradiated mice served as controls. (A) Lung and (B) heart tissues were harvested six months post-irradiation and stained with Masson’s Trichrome to assess fibrosis. Sample size: n = 5 mice per group. Data are presented as mean ± standard deviation. Statistical significance: p < 0.05, p < 0.01, p < 0.001, p < 0.0001.

**Figure S2.**
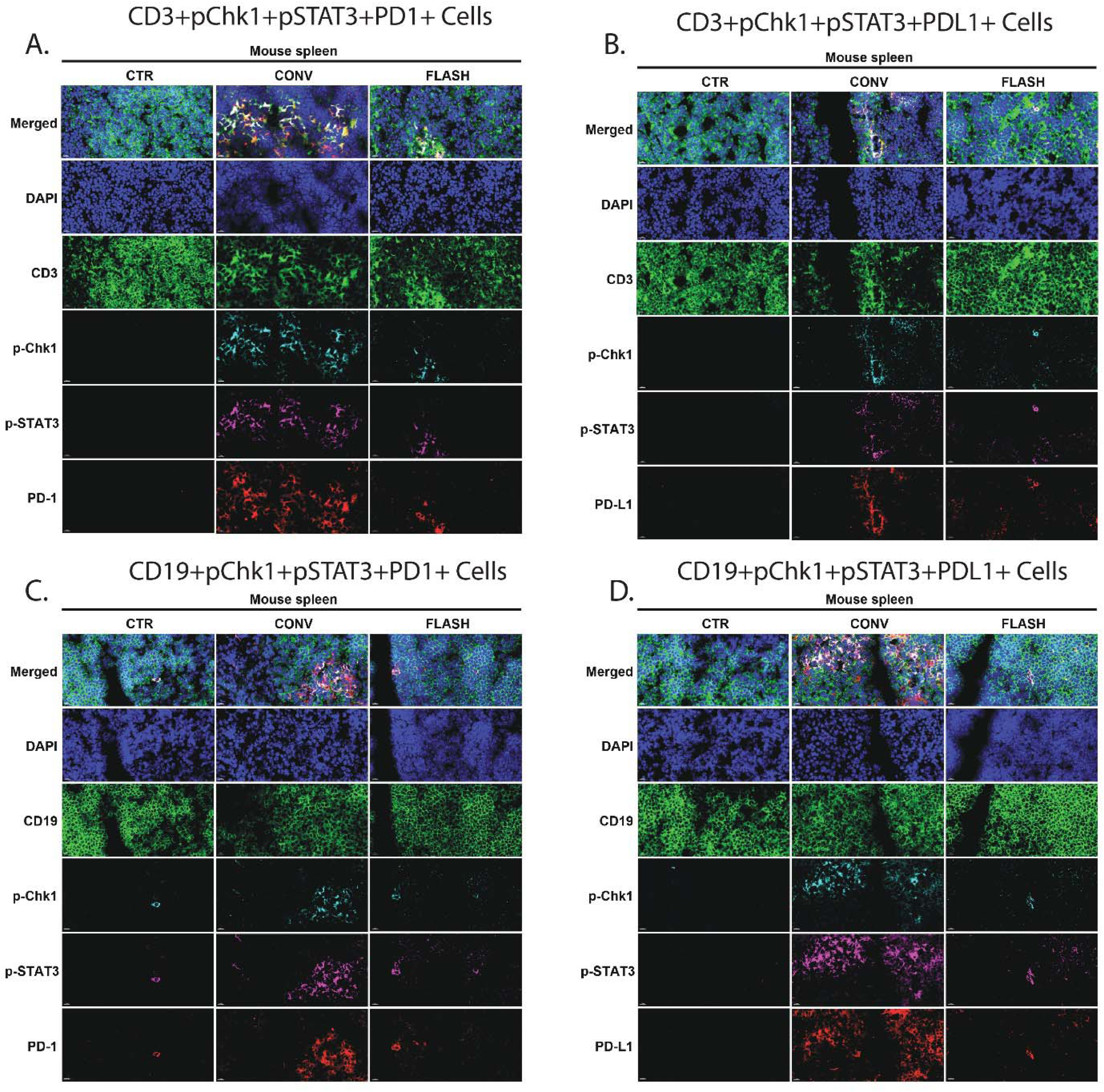
FLASH Radiotherapy limits Chk1-STAT3-mediated immune activation in lymphoid tissue. Spleens were harvested from eight-week-old C57BL/6 mice two months post-irradiation after receiving single-fraction (17 Gy) thoracic irradiation using either CONV or FLASH RT. Immunofluorescence staining was performed to assess expression of indicated markers. Scale bar, 10 µm. Representative images show co-expression of (A) CD3^⁺^p-Chk1^⁺^p-STAT3^⁺^PD-1^⁺^, (B) CD3^⁺^p-Chk1^⁺^p-STAT3^⁺^PD-L1^⁺^, (C) CD19^⁺^p-Chk1^⁺^p-STAT3^⁺^PD-1^⁺^, and (D) CD19^⁺^p-Chk1^⁺^p-STAT3^⁺^PD-L1^⁺^. Scale bar = 10 µm. Sample size: n = 5 mice per group.

**Table S1:** Physical Beam Parameters Measured using Gafchromic Film and Beam Current Transformers for Single Fraction and Five Fraction Irradiations.

| Treatment Group | Dose (Gy) | Number of Pulses (N) | Dose per Pulse (Gy) | Mean Dose Rate (Gy/s) | Irradiation Time (s) | Intrapulse Dose Rate (MGy/s) | Pulse Width (μs) | Pulse Repetition Frequency (PRF, Hz) | Fractions |
| --- | --- | --- | --- | --- | --- | --- | --- | --- | --- |
| FLASH | 17 | 7 | 2.43 | 340 | 0.05 | 1.4 | 1.9 | 120 | 1 |
| CONV | 17 | 1913 | 0.0088 | 0.3 | 57 | 0.0074 | 1.2 | 30 | 1 |
| FLASH | 2 | 1 | 2.00 | $1.4 \times 10^6$ | $1.4 \times 10^{-6}$ | 1.4 | 1.4 | NA | 5 |
| CONV | 2 | 225 | 0.0088 | 0.3 | 7 | 0.0074 | 1.2 | 30 | 5 |

## Notes

### Summary of Updates

Figures has been reorganized to reflect the order presented.

