## Supplementary figures and images for "FLASH Radiotherapy Mitigates Radiation-Induced Lymphopenia and Prevents Immunosuppression via Chk1-STAT3 Axis Modulation in a Preclinical Thoracic Irradiation Model"

### Figure S1

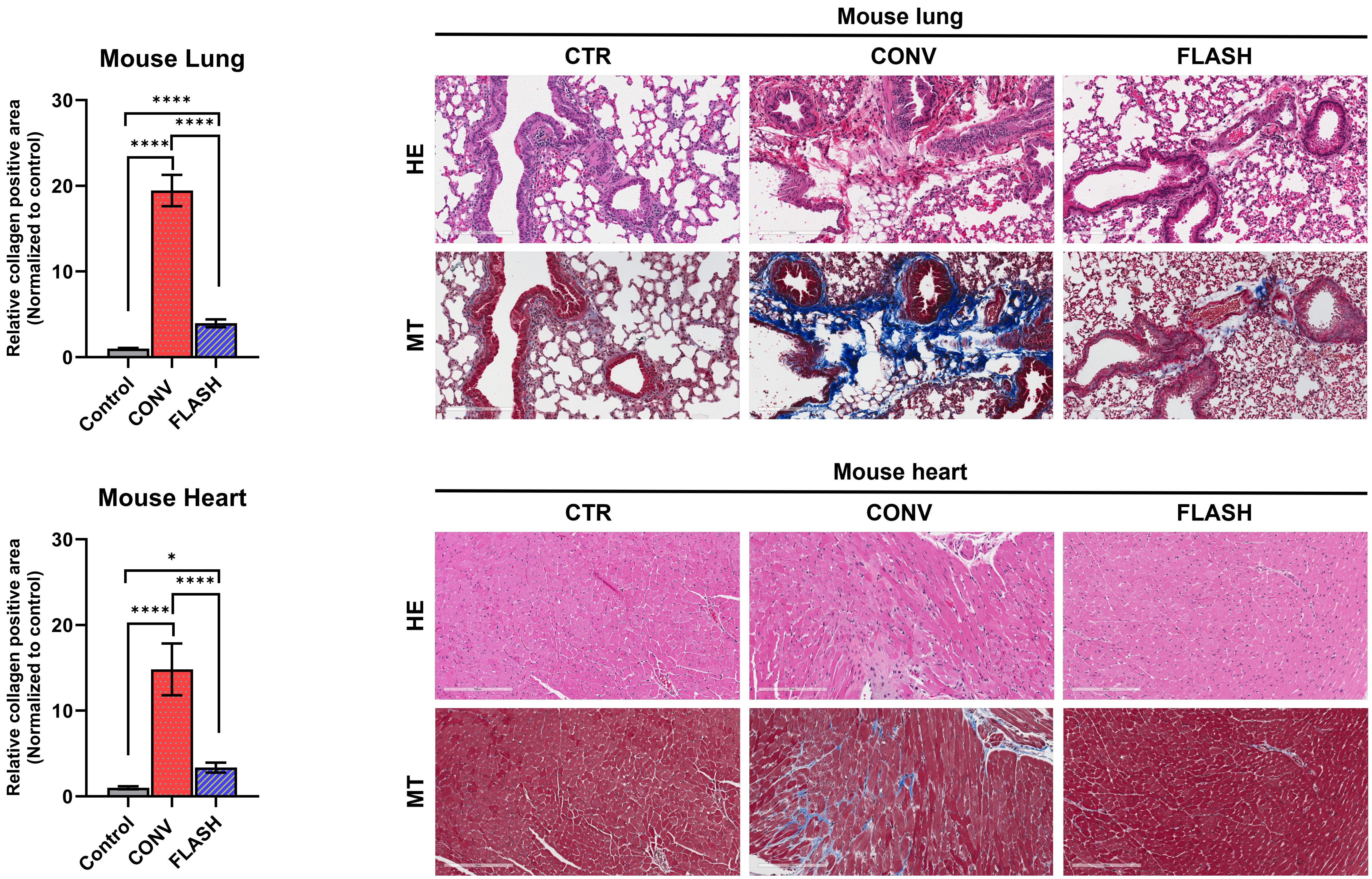

### Figure S2

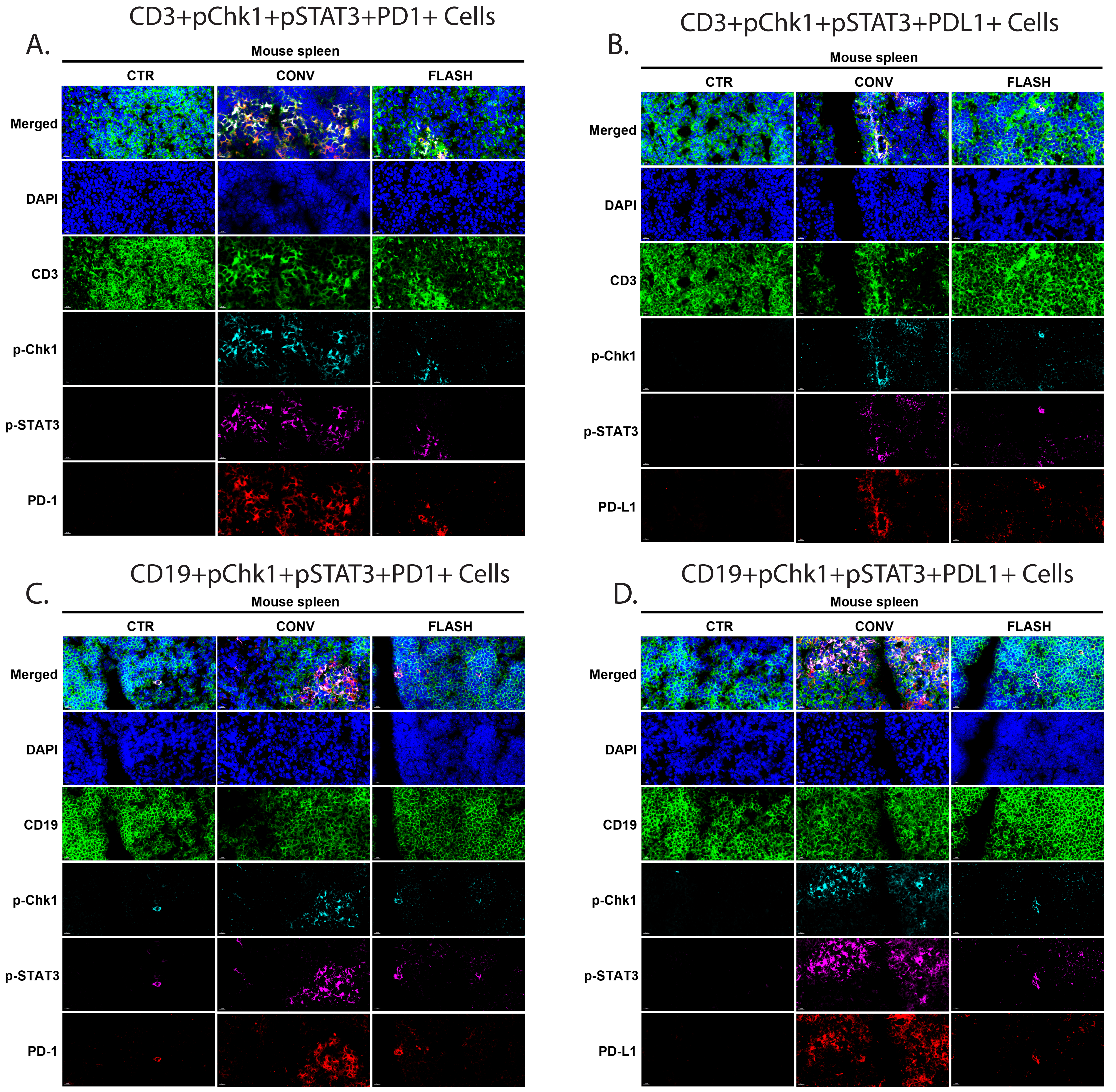
